## Supplemental information for "A divergent cyclic nucleotide binding protein promotes *Plasmodium* ookinete infection of the mosquito"

A

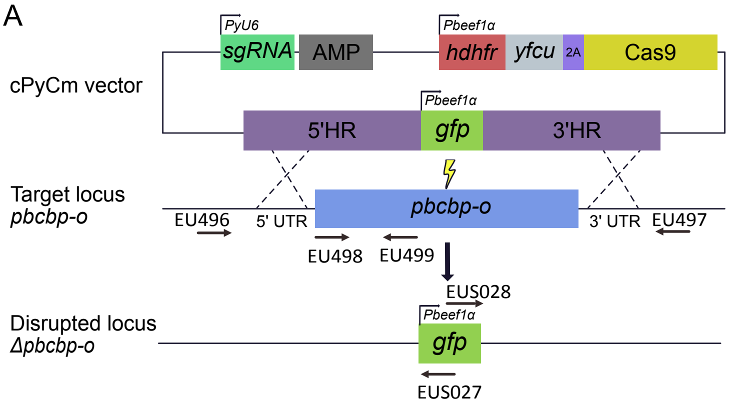


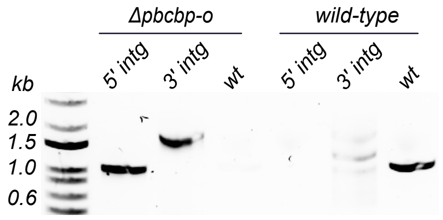


| Product | Primers | Size (kb) |
|---|---|---|
| 5' integration | EU496 + EUS027 | 1.1 |
| 3' integration | EU497 + EUS028 | 1.9 |
| wild type | EU498 + EU499 | 1.15 |

B

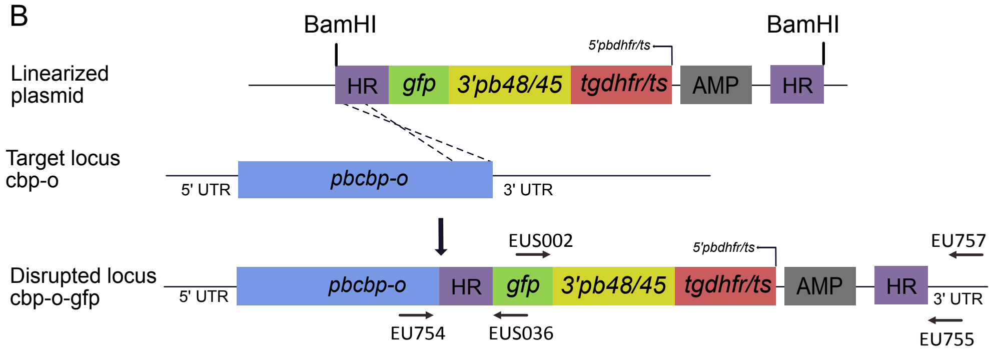


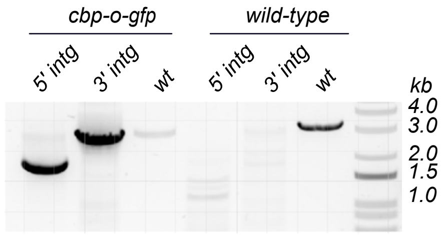


| Product | Primers | Size (kb) |
|---|---|---|
| 5' integration | EU754 + EUS036 | 1.7 |
| 3' integration | EUS002 + EU757 | 3 |
| wild type | EU754 + EU755 | 3 |

A

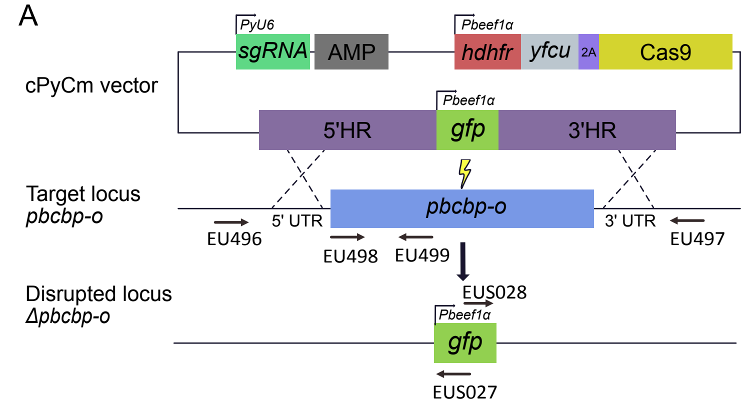


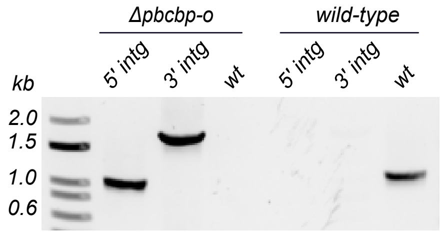


| Product | Primers | Size (kb) |
|---|---|---|
| 5' integration | EU496 + EUS027 | 1.1 |
| 3' integration | EU497 + EUS028 | 1.9 |
| wild type | EU498 + EU499 | 1.15 |

B

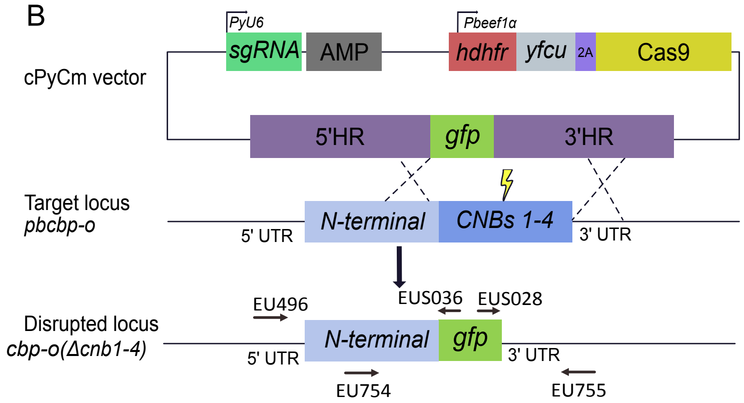


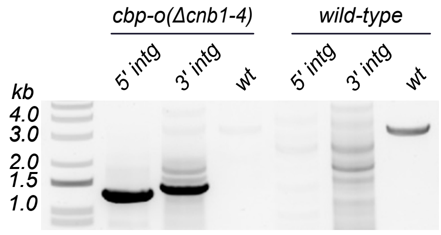


| Product | Primers | Size (kb) |
|---|---|---|
| 5' integration | EU756 + EUS036 | 1.2 |
| 3' integration | EU755 + EUS028 | 1.4 |
| wild type | EU755 + EU754 | 3 |

C

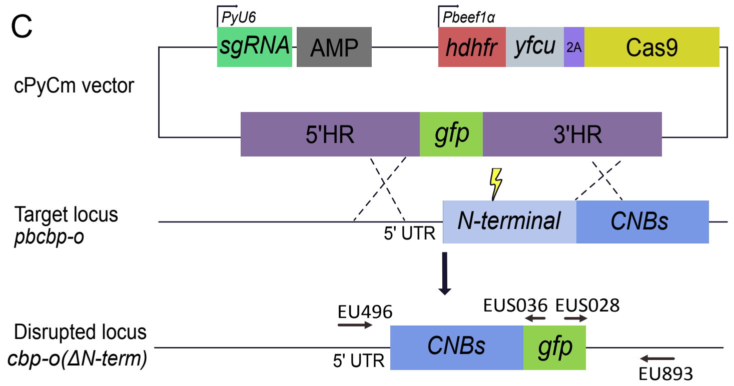


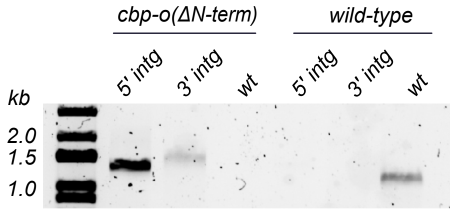


| Product | Primers | Size (kb) |
|---|---|---|
| 5' integration | EU496 + EUS036 | 1.2 |
| 3' integration | EUS028 + EU893 | 1.4 |
| wild type | EU498 + EU499 | 1.3 |

D

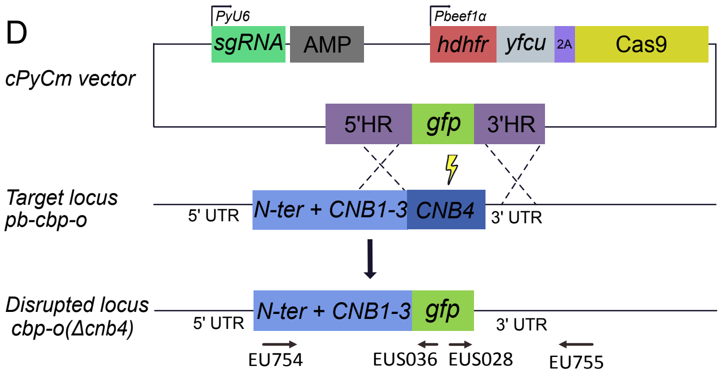


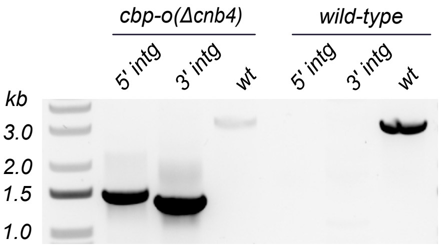


| Product | Primers | Size (kb) |
|---|---|---|
| 5' integration | EU754 + EUS036 | 1.5 |
| 3' integration | EUS028 + EU755 | 1.45 |
| wild type | EU754 + EU755 | 3.2 |

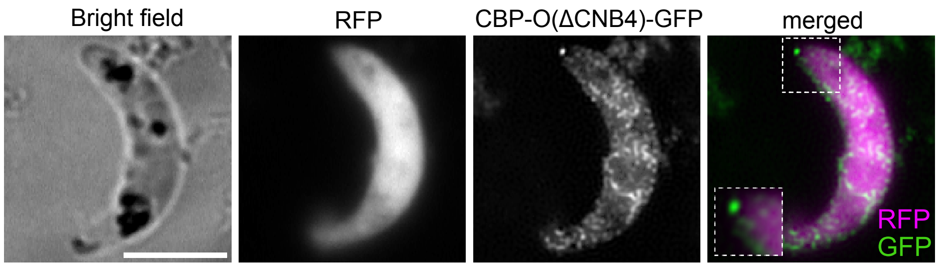
Bright field
RFP
CBP-O(ΔCNB4)-GFP
merged
RFP
GFP

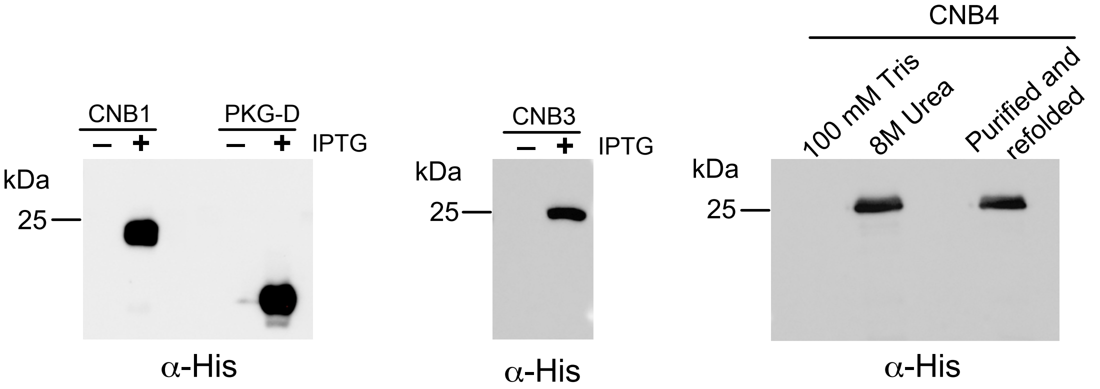
CNB1
PKG-D
– + – + IPTG
kDa
25
α-His
CNB3
– + IPTG
kDa
25
α-His
CNB4
100 mM Tris
8M Urea
Purified and refolded
kDa
25
α-His

**Supporting information**

SF1 Schematic representation of strategy to generate PbCBP-PO transgenic lines

(A) Full-length knock out was generated using CRISPR/Cas9 with a guide RNA mapping to the middle of the gene. The locus was replaced by a constitutively active GFP. (B) CBP-O was C- terminally tagged with GFP employing a single crossover strategy where the gene was linearised using a BamHI site. Positions of oligonucleotides used for genotyping are indicated by arrows and identified by specific EU numbers. Genotyping by the respective diagnostic PCRs and expected sizes are indicated by agarose gel electrophoresis and a table respectively.

SF2 Schematic representation of strategy to generate PbCBP-PO mutant lines

All mutant lines were created in constitutively expressing RFP line. (A) Full-length knock out (*Δcbp-o*) was generated using the strategy as represented in SF1A. (B) *cbp-o(Δcnb1-4),* (C) *cbp-o(ΔN-term)* and (D) *cbp-o(Δcnb4)* transgenic lines were generated by replacing the indicated loci with *gfp* resulting in a GFP-tagged domain mutant. Positions of oligonucleotides used for genotyping are indicated by arrows and denoted by specific EU numbers. Genotyping by diagnostic PCRs and expected sizes are indicated.

SF3. Deletion of CNB4 does not impact protein localisation to the apical tip.

Immunofluorescence microscopy of mature ookinete expressing GFP-tagged CBP-O protein lacking the terminal CNB4 domain.

SF4. Recombinant protein expression of PbCBP-O's cyclic nucleotide binding domains

Western blot analysis of His-tagged recombinant CNB1, CNB3 and CNB4 proteins. CNB1 and 3 were isolated from solution fractions after lysis under non-denaturing conditions. CNB4 was isolated from inclusion bodies in 8M Urea buffer, followed by refolding dialysis.

**Table 1: Sequences of primers used to confirm genomic integration**

| Primer | Sequence | Name |
|---|---|---|
| EU496 | TTCCCTTGTCTTTGTAGCCCATAG | 5’ ΔCBP-O<br>5’ CBP-O(ΔN-term) |
| EUS027 | GCAATTAATGTGAGTTAGCTC | 5’ ΔCBP-O |
| EU497 | GGTTACGCACCCAAAATTGTTG | 3’ ΔCBP-O |
| EUS028 | GCTGCTGGGATTACACATGG | 3’ ΔCBP-O<br>3’ ΔCBP-O(ΔN-term)<br>3’ CBP-O(ΔCNB4)<br>3’ CBP-O(ΔCNB1-4) |
| EU498 | GGCCAAGATGTCTGCTATTTCAAAG | wt ΔCBP-O<br>wt ΔCBP-O(ΔN-term) |
| EU499 | GTACATGCTTTCTCTTTGTTCGTTGG | wt ΔCBP-O<br>wt ΔCBP-O(ΔN-term) |
| EU754 | TGGGTTGAAAGAAGCACACAATG | wt/ 5' CBP-O-GFP<br>wt CBP-O(ΔCNB1-4)<br>wt/ 5’ CBP-O(ΔCNB4) |
| EUS036 | GTATCTCGCAAAGCATTGAACACC | 5' CBP-GFP<br>5' CBP-O-(ΔCNB1-4)<br>5' CBP-O(ΔN-term)<br>5’ CBP-O(ΔCNB4) |
| EUS002 | GATTAAGTTGGGTAACGCCAG | 3' CBP-O-GFP |
| EU755 | GATTAAAAGTGAGGGTTACGCACC | wt CBP-O-GFP<br>3'CBP-O(ΔCNB1-4)<br>wt/ 3’ CBP-O(ΔCNB1-4)<br>wt/ 3’ CBP-O(ΔCNB4) |
| EU757 | GTTACGCACCCAAAATTGTTG | 3' CBP-O-GFP |
| EU756 | GAAGGATCGGAGTGCCCATATAC | 5' CBP-O(ΔCNB1-4) |
| EUS028 | GCTGCTGGGATTACACATGG | 3' CBP-O(ΔCNB1-4)<br>3' CBP-O(ΔN-term) |
| EU893 | TGGATATACTCTTTCCTCCATTATCCG | 3' CBP-O(ΔN-term) |

**Table 2 Sequences of primers for plasmid constructs**

| | | |
|---|---|---|
| EU492 | aatGGTACCTTATGAAGGAATAGCAAGAG | ΔCBP-O 3'HR |
| EU493 | aatGCTAGCTATAGCATGTGCTAATTCTTTAAAC | ΔCBP-O 3'HR |
| EU503 | ccgAAGCTTCAATGAAATATTTGTACGCAATG | ΔCBP-O 5'HR |
| EU504 | aatCCGCGGTTCAATATACACATATACCTTGTC | ΔCBP-O 5'HR |
| EU564 | CATACTTCGAGTTATACAATGTTATTGTTTCTAGAAA<br>TAACAAACACGTTTTAGAGCTAGAAATAGCAAGTT | ΔCBP-O gRNA |
| EU700 | CATACTTCGAGTTATACAATGTTATTGCTTCGAACA<br>AATCAAGTTAAGTTTTAGAGCTAGAAATAGCAAGTT | CBP-O(ΔCNB1-4)<br>gRNA |
| EU749 | CTATGACCATGATTACGCCAAGCTTGATTTAGGCAT<br>TGAGTTAAATGAAG | CBP-O(ΔCNB1-4)<br>5'HR |
| EU750 | CATTGAAGACCGCGGAGGTATATTATCAACAATACT<br>CATAAATTC | CBP-O(ΔCNB1-4)<br>5'HR |
| EU751 | TAATATACCTCCGCGGTCTTCAATGAGTAAAG | CBP-O(ΔCNB1-4)<br>3'HR |
| EU752 | ATTAAATTGTAAACTTAAGGAATTCTATAGCATGTGC<br>TAATTCTTTAAACAAG | CBP-O(ΔCNB1-4)<br>3'HR |
| EU708 | aatCCGCGGTCTTCAATGAGTAAAGGAGAAGAACT<br>TTTC | CBP-O(ΔCNB1-4)<br>GFP |
| EU709 | ccgcGGTACCTTATTTGTATAGTTCATCCATGCC | CBP-O(ΔCNB1-4)<br>GFP |

| | | |
|---|---|---|
| EU883 | catacttcgagttatacaatgttattGCATGTCTAGAATGTCCCAAAgttttagagctagaaatagcaagtt | CBP-O(ΔN-term) gRNA |
| EU884 | ctatgaccatgattacgccaagcttCAATGAAATATTTGTACGCAATGC | CBP-O(ΔN-term) 5'HR |
| EU885 | gctccgcggctcctttactcatattatttttataGACATTGTATAAATCGTCGTTTTTTATATC | CBP-O(ΔN-term) 5'HR |
| EU886 | ggggcggccgccaatgtctataaaaataatATGAGTAAAGGAGAAGAACTTTTCAC | CBP-O(ΔN-term) GFP |
| EU887 | tgagctgaagaccgcggTTTGTATAGTTCATCCATGCCATG | CBP-O(ΔN-term) GFP |
| EU888 | tatacaaaccgcggtcttcaGCTCATACAAATGTTGCTATAGTCG | CBP-O(ΔN-term) 3'HR |
| EU889 | gggggtaccattaaattgtaaacttaaggaattcTGACGTATTGAAACTACACCTTG | CBP-O(ΔN-term) 3'HR |
| EU706 | acgTCTAGAGAACTAAACTTAAGGTTATGTGAAATG | CBP-O-GFP |
| EU707 | aatCCATGGaTGTATTAATTTTATTGTCAGGGTTTTG | CBP-O-GFP |
| EU701 | CATACTTCGAGTTATACAATGTTATTGTCGTATCCTCGGAATTTAATGTTTTAGAGCTAGAAATAGCAAGTT | CBP-O(ΔCNB4) gRNA |
| EU704 | gccgcAAGCTTAACAGTTGATGATGCTATTATAAAC | CBP-O(ΔCNB4) 5'HR |
| EU705 | aatCCGCGGTGGGACATTTTTAAGTACTTG | CBP-O(ΔCNB4) 5'HR |
| EU831 | GGAGTGCCATATGGATGCATCTATAGATTATAATAATAAAAAAAGTATC | for-PbPKG-D |
| EU832 | GATACTCGAGtcaAAATGTACCTCTCCCTATTATTC | rev-PbPKG-D |
| EU839 | CGAAGCTAGCGCTCATACAAATGTTGCTATAGTCG | for-CNB1 |
| EU840 | GCATCCTCGAGtcaCATATTCAAACGGGATATTGATATGTTTATGTC | rev-CNB1 |
| EU835 | GCAACTCATATGATACAAAAAAAAGATGGAGACTTTATAG | for-CNB3 |
| EU836 | CATACTCGAGttaATAGTTATTTTCATTACCATATATGGTTTTTAG | rev-CNB3 |
| EU837 | CGAACCTCATATGATGTTAGAAGAAAGACAACAAAATTG | for-CNB4 |
| EU838 | CCATCTCGAGttaTGTATTAATTTTATTGTCAGGGTTTTG | rev-CNB4 |
